# CD11c-expressing microglia are transient, driven by interactions with apoptotic cells

**DOI:** 10.1101/2024.06.24.600082

**Authors:** Nathaniel Ghena, Sarah R Anderson, Jacqueline M Roberts, Emmalyn Irvin, Joon Schwakopf, Alejandra Bosco, Monica L Vetter

## Abstract

Microglia, the parenchymal macrophage of the central nervous system serve crucial remodeling functions throughout development. Microglia are transcriptionally heterogenous, suggesting that distinct microglial states confer discrete roles. Currently, little is known about how dynamic these states are, the cues that promote them, or how they impact microglial function. In the developing retina, we previously found a significant proportion of microglia express CD11c (Integrin αX, complement receptor 4, *Itgax*) which has also been reported in other developmental and disease contexts. Here, we sought to understand the regulation and function of CD11c+ microglia. We found that CD11c+ microglia track with prominent waves of neuronal apoptosis in postnatal retina. Using genetic fate mapping, we provide evidence that microglia transition out of the CD11c state to return to homeostasis. We show that CD11c+ microglia have elevated lysosomal content and contribute to the clearance of apoptotic neurons, and found that acquisition of CD11c expression is, in part, dependent upon the TAM receptor Axl. Using selective ablation, we found CD11c+ microglia are not uniquely critical for phagocytic clearance of apoptotic cells. Together, our data suggest CD11c+ microglia are a transient state induced by developmental apoptosis rather than a specialized subset mediating phagocytic elimination.

## Introduction

Microglia, the parenchymal macrophage of the central nervous system (CNS), serve crucial remodeling functions throughout development, including the refinement of circuits and cell elimination (1, 2). Single-cell analysis has revealed that microglia are transcriptionally heterogeneous during development (3, 4). However, the cues that produce specific states, the functional role(s) associated with these states, and how dynamic and reversible these states are not well understood.

The developing retina is an ideal CNS system to link well-characterized developmental events with microglial properties and function. We previously found that murine retinal microglia gene expression is heterogeneous and dynamic throughout development (5, 6). Notably, we showed that a majority of microglia early in postnatal retina express CD11c (Integrin αX, complement receptor 4, *Itgax*) and were remarkably similar to CD11c-expressing (CD11c+) microglia of the developing and diseased brain (6, 7). By single-cell RNA sequencing, we found heterogeneity in CD11c+ microglia, but all CD11c+ microglia expressed genes associated with phagocytosis and lysosomal function (6). In the developing CNS, CD11c+ microglia predominantly reside in white matter tracts (8-10) and have dynamic regional density changes over development in brain and spinal cord (11). CD11c+ microglia are increased in diverse disease states. CD11c is a marker of disease-associated microglia (DAM, MgND), enriched around Aβ plaques in Alzheimer’s disease (12, 13). The appearance of CD11c+ microglia in these diverse contexts raises questions regarding their regulation and function. To address this, we sought to understand the developmental cues and roles of CD11c+ microglia in developing retina.

We previously showed a link between apoptosis and CD11c-expression, as *in vivo* loss of developmental apoptosis dramatically reduced the proportion of CD11c+ retinal microglia and expression of phagocytic genes (6). Consistent with this, microglia upregulate CD11c and coexpress Osteopontin (OPN) in response to the engulfment of apoptotic neurons in cell culture (14). In addition, CD11c+ microglia within developing white matter tracts engulf oligodendrocytes (9, 15), suggesting that CD11c+ microglia may play a key role in phagocytosis. We found that the proportion of CD11c+ microglia over retina development is dynamic, which raises the question of whether developmental CD11c+ microglia represent a transient state associated with cell death and phagocytic function, and whether these cells mediate a specialized role in engulfment.

Here, we find that CD11c+ microglia density in the retina coincides with waves of neuronal apoptosis, that CD11c+ microglia predominately associate with dying neurons and are phagocytic. Through genetic fate mapping, we find that CD11c+ microglia shift to CD11c-state after periods of developmental apoptosis. Additionally, we find that loss of Axl reduced the proportion of CD11c+ cells, implicating Axl-mediated recognition of apoptotic cells in driving CD11c+ microglial state. Using selective depletion, we find both CD11c+ and CD11c-microglia contribute to the timely clearance of apoptotic neurons in the retina. Altogether, these data suggest that CD11c+ microglia in the developing nervous system represent a dynamic population driven by phagocytic interactions with apoptotic cells.

## Results

### CD11c-microglia coincide with developmental cell death

We previously found an increased proportion of CD11c+ microglia in postnatal retina compared to embryonic or adult stages, partially dependent on developmental cell death (6). We therefore hypothesized that CD11c+ microglia arise in response to interactions with apoptotic cells. To test this, we first utilized CD11c-DTR/GFP transgenic mice, in which the *Itgax* promoter (CD11c) directs expression of a diphtheria toxin receptor-enhanced green fluorescent protein fusion protein (DTR/GFP) (16). We confirmed via flow cytometry that GFP expression accurately correlates with surface expression of CD11c protein in naïve CD11c-DTR/GFP mouse retinas (Supplementary Figure 1A). Next, we quantified the density of CD11c+ cells by GFP expression and found the density of CD11c+ cells peaks between postnatal (P) 5 and P10, and drops by P30 (Figures 1A, B). Using complement component 1q (C1q), which specifically and stably labels microglia in the developing retina (5), we confirmed CD11c-GFP+ cells are microglia and peak at P5 consistent with our previous single-cell sequencing data (Figure 1C, D; (6)). Morphologically, CD11c+ microglia become more complex over the course of development, similar to non-CD11c microglia (Figure 1E; (6)). By analyzing retinal cross-sections, we found that CD11c-GFP+ microglia distribute primarily in the ganglion cell layer (GCL) until P3, followed by a shift to the inner nuclear layer (INL) at P7, spatiotemporally tracking with developmental cell death (Figures 1F, G; (17, 18)). Altogether, CD11c+ microglia track with periods of cell death across development and within specific retinal subregions.

**Figure 1.**
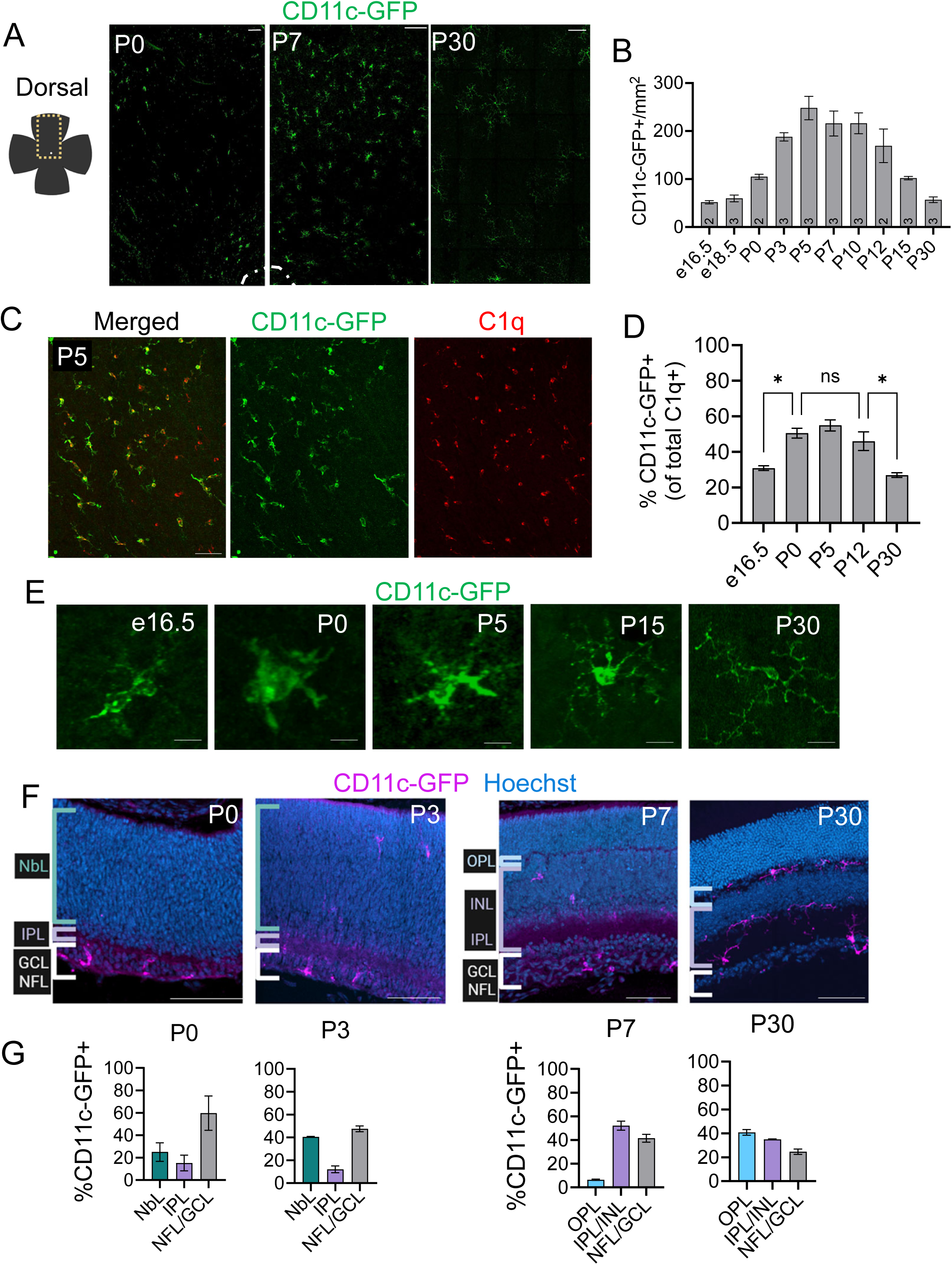
CD11c+ microglia density and localization track with periods of apoptosis. (A) *Left* – schematic of retinal flat mount depicting where analysis was conducted, with dorsal leaf outlined in dashed yellow box. *Right* – max projected confocal images of CD11c-GFP+ cells in the dorsal leaf of retinal flat mounts at postnatal days (P) 0, 7, and 30. Optic nerve head outlined in dashed white line. Scale bar, 100μm. (B) Densities of CD11c-GFP+ cells from retinal flat mount across embryonic and postnatal ages. Data are presented as means ± SEM (2-3 animals age). One-way ANOVA F(9,17) =18.87 P<.0001. (C) Representative Max projected confocal image of C1q+ (red) and CD11c-GFP+ (green) microglia at P5. Scale bar 50μm. (D) Proportion of CD11c-GFP+ retinal microglia from retinal flat mounts at embryonic day (e) 16.6, P0, P5, P12, and P30. Data are presented as mean ± SEM (2-3 animals/age. One-way ANOVA F(4, 5) =15.76 *P=.0049* and Tukey’s multiple comparisons. \**P* ≤.05, *NS* =no significance. (E) Max projected confocal images of individual CD11c-GFP+ microglia in retinal flat mounts from embryonic day 16.5 (e16.5) through postnatal day 30 (P30). Scale bar at 10μm. (F) Confocal images of retinal cross sections from CD11c-DTR/GFP transgenic mice at P0, P3, P7, and P30. CD11c-GFP (Magenta); Hoechst (blue). For P0/3, brackets segment retinal layers, gray – nerve fiber layer and ganglion cell layer (NFL/GCL), purple – inner plexiform layer (IPL), and teal – neuroblastic layer (NbL). P7/P30, gray – nerve fiber layer and ganglion cell layer (NFL/GCL), purple – inner plexiform layer and inner nuclear layer (IPL/INL), and blue – outer plexiform layer (OPL) for P7 and P30. Scale bar, 100μm. (G) Percentage of CD11c-GFP+ microglia across all retinal layers at P0, P3, P7, and P30 (n = 2 animals). Data are presented as means ± SEM.

### Microglia transition out of the CD11c state

Since we found the proportion of CD11c-GFP+ microglia shifts throughout retina development, we predict CD11c+ microglia are a transient state rather than a stable population. To test this, we performed lineage-tracing to identify microglia that have expressed CD11c by utilizing CD11c-Cre/GFP transgenic mice (19) crossed with Rosa-Ai14 tdTomato reporter mice (20) (Figure 2A). With this strategy, cells actively expressing CD11c will express GFP while cre-mediated recombination stably marks cells that previously expressed CD11c (Figure 2A). Therefore, cells expressing CD11c (CD11c-active) would be GFP^+^ tdTomato^+^ or GFP^+^tdTomato^-^, while cells that previously expressed CD11c (CD11c-lineage) would be GFP^-^ tdTomato^+^, and cells that never expressed CD11c (CD11c-negative) would be GFP^-^ tdTomato^-^ (Figure 2A). Using flow cytometry, we found that the proportion of microglia CD11c-active matched our previous quantification of CD11c+ microglia at P5 and P30 (∼50% and <20%, respectively) while surprisingly, the majority of the remaining microglia were CD11c-lineage. This suggests that nearly all postnatal retinal microglia have either previously express or are currently expressing CD11c (Figure 1E; Figure 2B). Interestingly, the proportion of cells marked as CD11c-lineage increased from P5 to P30 in a manner proportional to the reduction of CD11c-active cells (Figure 2B), further evidence that CD11c-expression is transient and CD11c+ microglia convert to non-CD11c microglia. The P5 CD11c-active microglia population appeared more phagocytic than the CD11c-lineage microglia at P5 as they had higher CD68 area, were more likely to have phagocytic cups, and had more cups per microglia (Figure 2C – F). Bulk RNA-sequencing on sorted P5 retinal microglia comparing CD11c-active and CD11c-lineage microglia confirmed that CD11c-active have higher expression of genes associated with phagocytosis and remodeling states *(Lamp1, Lyz2, CD68*) while CD11c-lineage have higher expression of homeostatic genes (*P2ry12, Tgfbr1, Tmem119*) (Figure 2G). Altogether, these findings suggest that CD11c+ microglia are more phagocytic and are not a stable subset, but shift to a non-CD11c state after resolution of periods of developmental apoptosis.

**Figure 2.**
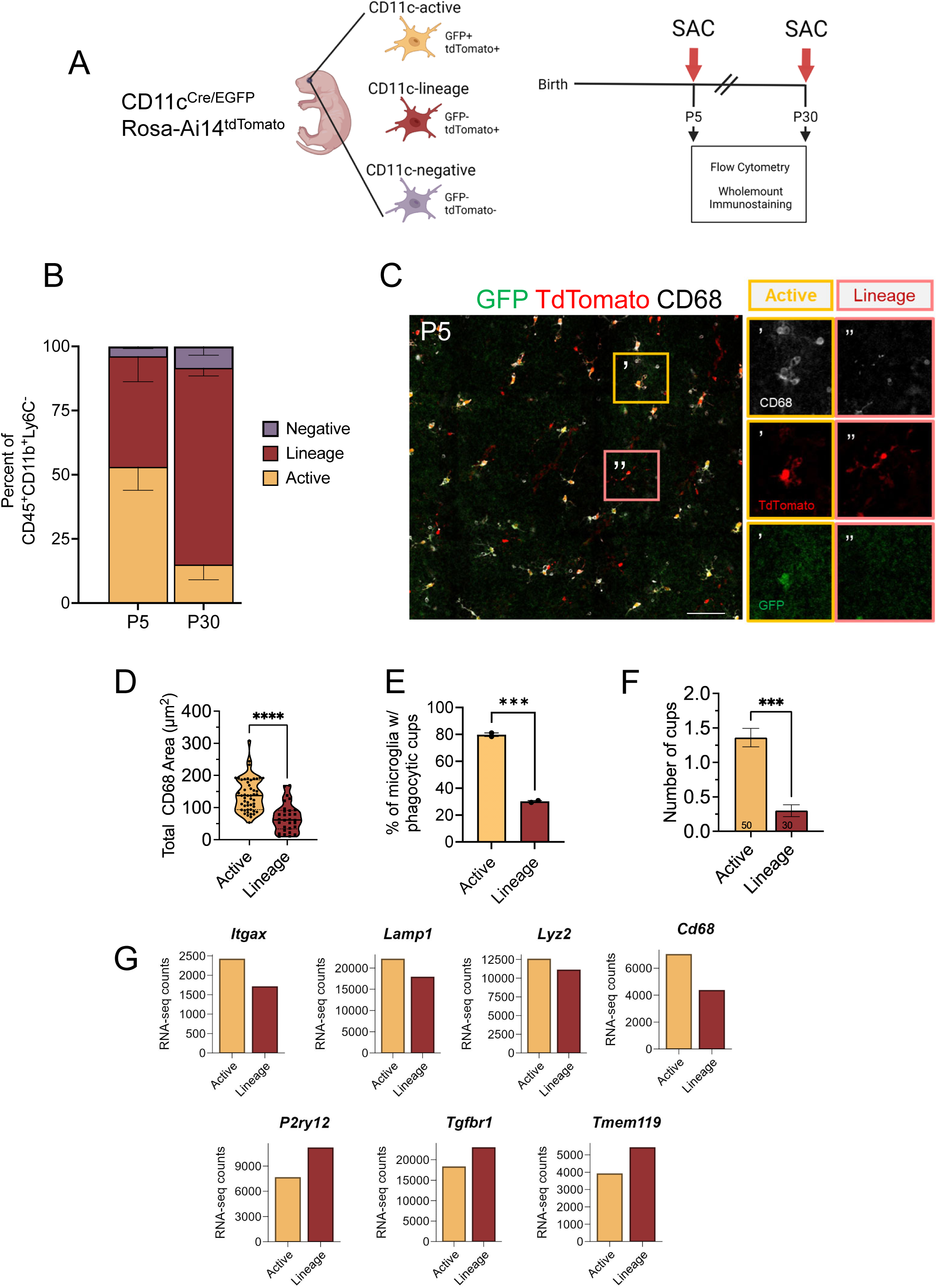
CD11c+ microglia are a dynamic state. (A) Cartoon depiction of CD11c lineage tracing strategy crossing CD11c-Cre/EGFP mice with Rosa-Ai14 tdTomato and the analysis performed at P5 and P30. (B) The proportion of microglia at P5 and P30, comparing microglia that are CD11c-active (GFP+tdTomato+/total microglia), CD11c-lineage (GFP-tdTomato+/total microglia), and CD11-negative (GFP-tdTomato-/total microglia) by flow cytometry. (C) Max Projected confocal image from P5 CD11c-Cre^EGFP/+^; Rosa^tdTomato/+^ retinal flat mount. Insets ‘ CD11c-Active (gold box) and “ CD11c-Lineage (pink box) microglia. CD68 (white), CD11c-Active (GFP, green), CD11c-Lineage (tdTomato, red). Scale bar, 50μm. (D) Quantification of CD68 area in CD11c-Active and CD11c-Lineage microglia at P5 (n=2 animals, 30 CD11c-Lineage cells and 50 CD11c-Active cells) ± SEM. Mann-Whitney test \*\*\*\**P* <.0001. (E) The proportion of CD11c-active and CD11c-lineage microglia with phagocytic cups at P5 (n=2 animal) ± SEM. Unpaired t test *\*\*\*P* <.001. (F) Quantification of number of phagocytic cups in CD11c-Active and CD11c-Lineage microglia at P5 (n=2 animal, 30 CD11c-Lineage cells and 50 CD11c-Active cells) ± SEM. Unpaired t-test *\*\*\*P* <.001. (G) Bar graphs of RNA-seq read counts for (top) genes with increased counts in CD11c-active and (bottom) genes with decreased counts in CD11c-lineage.

### CD11c-microglia phagocytose apoptotic cells and are regulated by Axl signaling

Since we observed CD11c-expression in microglia to be transient, coinciding with periods of developmental apoptosis, we asked whether CD11c+ microglia interacted with dying neurons *in vivo.* At P3, during peak retinal ganglion cell (RGC) developmental apoptosis, we found CD11c-GFP+ microglia interacted with and engulfed apoptotic RGCs, as co-labeled by activated caspase 3 (CC3+) and RGC marker RNA-binding protein with multiple splicing (RBPMS), with most CD11c-GFP+ microglia directly interacting with RBPMS+CC3+ cells (∼70%; Figures 3A, B). Immunostaining for lysosomal marker CD68 and microglial marker C1q, we found that the higher CD11c-GFP expression correlated with elevated levels of CD68 and an enlarged amoeboid morphology, consistent with increased lysosomal content and phagocytic/digestion activity (Figure 3C – E). This suggests that CD11c-expression is driven by recent interactions and/or engulfment of apoptotic cell bodies.

**Figure 3.**
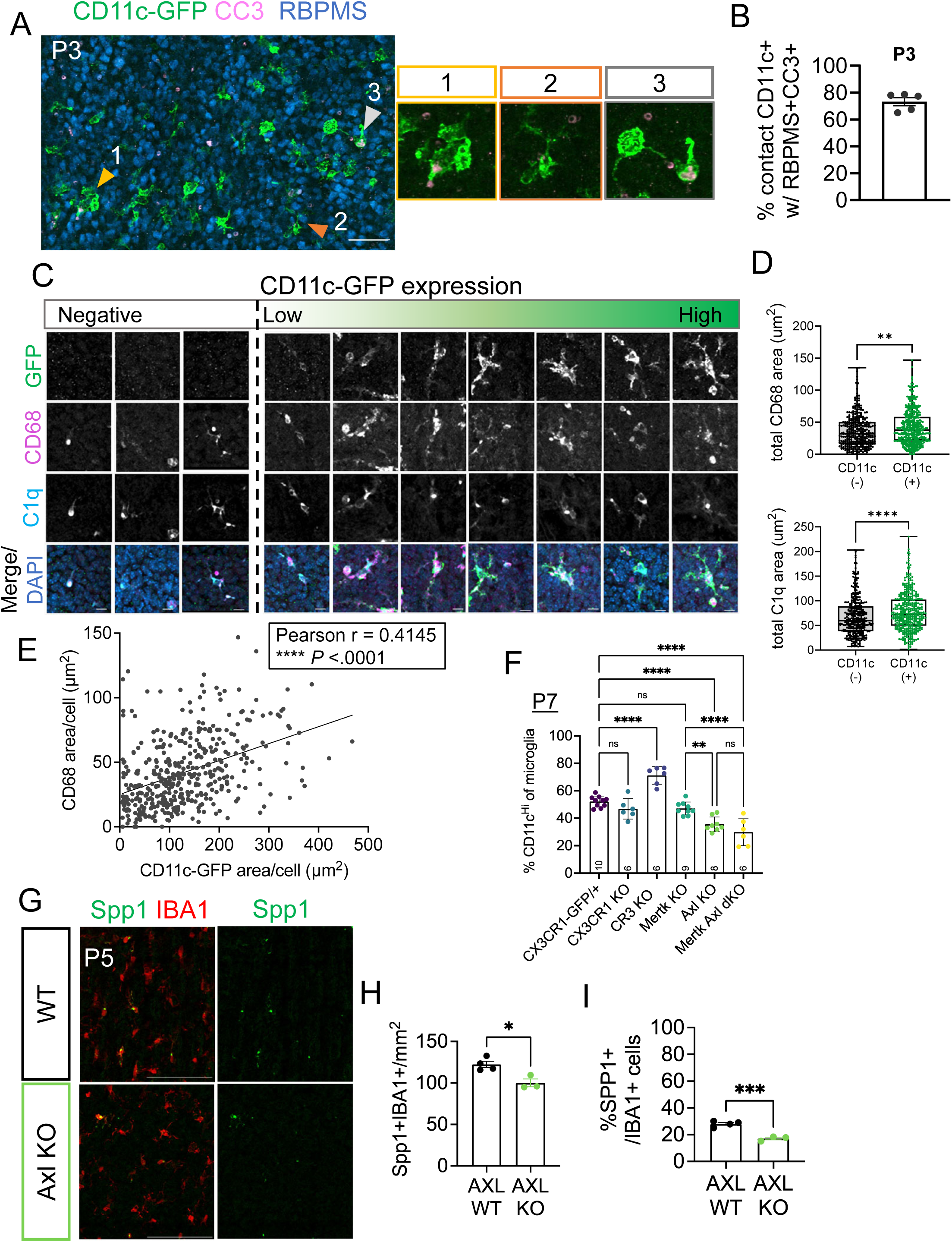
CD11c+ microglia emerge from interactions with apoptotic cell bodies, mediated in part by Axl signaling. (A) Confocal image of P3 retinal flat mount (Arrowhead and magnified boxes illustrate interactions) in NFL/GCL. CD11c-GFP+ microglia (green); cleaved caspase 3, CC3 (pink); and RBPMS (blue). Scale bar, 50μm. (B) Percent CD11c-GFP+ microglia contacting apoptotic RGCs (RBPMS+CC3+) at P3 out of total CD11c-GFP+ microglia within GCL. (n=5 animals). (C) Confocal images of retinal flat mounts of P3 CD11c-GFP animals in GCL showing a range of microglial GFP expression from none to high. Scale bar, 10μm. Dashed line demarcates distinction between CD11c-GFP negative and positive. (D) Cell area immunostained for CD68 (*Top*) and C1q (*Bottom*) in CD11c-GFP- and CD11c-GFP+ microglia (n=4 animals, 352 GFP-cells and 422 GFP+ cells). Mann-Whitney test *Top **P =.*0011 and *Bottom ****P <.*0001. (E) Scatter plot of CD68 area/cell compared to CD11c-GFP area/cell at P3 in GCL. Pearson r= 0.4145 \*\*\*\**P* < .0001. (F) Flow cytometry analysis showing the percent CD11c^Hi^ of total CD45+CD11b+ or CD45+CX3CR1-GFP+ microglia from retinas across all genotypes. CX3CR1-GFP/+ (n=10), CX3CR1-KO (n=6), CR3 KO (n=6), MerTK KO (n=9), Axl KO (n=8), and MerTK/Axl dKO (n=6), ≥ 2 litters collected for each genotype, ± SEM. One-way ANOVA F(5, 39) = 34.64 *P*<.0001 and Tukey’s multiple comparisons. \*\*\*\**P*<.0001, \*\*\**P*=.0003, \*\**P*=.0056, \**P*=.02, *NS,* not significant. (G) Max projected confocal image of P5 retinal flat mount from WT and AXL KO, Spp1+ (green) and total microglia (IBA1+; red). Scale bar, 100μm. (H) Density of Spp1+IBA1+ microglia in AXL WT and KO (n=5, n=4 animals) ± SEM. Unpaired t test \**P* = .0139. (I) Percent Spp1+IBA1+ of total IBA1+ microglia in AXL WT and KO (n=5, n=4 animals) ± SEM. Unpaired t test *\*\*\*P* <.001

Since exposure to apoptotic neurons increases CD11c-expression (6, 14, 21), we asked which recognition pathways may be involved. Using flow cytometry, we analyzed genetic knockouts (KOs) of receptors previously implicated in the recognition and phagocytosis of apoptotic neurons, namely fractalkine receptor (CX3CR1, (22)), integrin receptor complement receptor 3 (CD11b/CR3, (23)), and TAM receptors Mer and Axl (24, 25) (Figure 3F, Supplementary Figure 2A – F). While we previously found that apoptotic RGC clearance was dependent on CR3 and Mertk (5), here we found that the proportion of CD11c+ microglia was significantly reduced in both Axl KO and Mertk/Axl double KOs, with approximately half the percentage of retinal microglia expressing CD11c compared to CX3CR1^GFP/+^ controls. Therefore, Axl regulates the acquisition of CD11c expression by microglia (Figure 3F). Additionally, we found an increase in CD11c-expression with loss of CR3, though this is likely due to a compensatory effect (26-28). Previously, we found Osteopontin (Spp1) RNA expression in microglia to be downregulated with loss of Axl (5), and CD11c-expression has been linked to Spp1-expression in various contexts (14, 29). Therefore, we assessed changes in Spp1 protein expression following loss of Axl signaling. We found the proportion and density of microglia expressing Spp1 is significantly reduced compared to wildtype controls (Figure 3G – I). Taken together, our results suggest CD11c+ microglia interact with apoptotic neurons, have increased phagocytic activity, and are partially regulated by Axl signaling.

### CD11c+ microglia are not specifically required for phagocytic clearance of apoptotic neurons

Next, we sought to determine whether CD11c+ microglia were specifically dedicated to the phagocytic clearance of dying cells. We utilized two depletion strategies to target CD11c+ and non-CD11c subsets (Figure 4A). First, to selectively ablate CD11c-GFP+ cells, we administered diphtheria toxin (DT) to CD11c-DTR/GFP pups at P0, P2, and P4, and collected retinas at P5. Primate diphtheria toxin receptor (DTR) expressed under the CD11c promoter allow for targeted ablation of CD11c-GFP+ cells using DT (30). We confirmed that DT administration did not affect microglia lacking expression of DTR as there was no change in total microglia density (C1q+/CX3CR1-GFP+) in CX3CR1-GFP mice following the same DT administration paradigm (Supplemental Figure 3A, B). DT administration to CD11c-DTR/GFP pups resulted in an ablation of ∼75% of CD11c-GFP+ microglia, constituting a 25% reduction in the total microglia pool at P5 (Figures 4B – E). Conversely, since we previously found that inhibition of CSF1R signaling eliminated homeostatic microglia while enriching for CD11c+ microglia (6), for our second depletion paradigm we administered CSF1R inhibitor pexidartinib (PLX 3397; (31)) to CX3CR1-GFP pups at P2 through P4, and collected at P5 (Figure 4A). As previously reported, we found the density of total C1q+ microglia was reduced by approximately 50% in PLX3397 treated animals (Figures 4F, G). Confirming our previous findings that CD11c-GFP+ microglia are less sensitive to CSF1R inhibition (5, 6), we found PLX3397 treatment resulted in virtually no reduction in CD11c+ microglia density, with the proportion of CD11c-GFP+ microglia shifting from ∼55% to ∼85% (Supplementary Figure 3C – F). To determine whether targeting these different microglia subsets had distinct effects on the clearance of apoptotic cells, we analyzed the density of dying RGCs as assessed by persistence of cleaved caspase 3 labeled RGCs (CC3+RBPMS+; Figure 4H). We found that DT-mediated depletion of CD11c-GFP+ microglia significantly increased the density of apoptotic RGCs compared to both Vehicle and WT+DT controls (Figure 4I). We showed the DT administration did not affect total RBPMS+ RGC number, confirming DT did not induce RGC death (Supplemental Figure 3G, H). We found that PLX3397-mediated microglia depletion also resulted in a significant persistence of apoptotic RGCs (Figures 4J). To examine whether the buildup of apoptotic RGCs was proportional to microglia density regardless of depletion strategy, we plotted CC3+RBPMS+ RGC density versus C1q+ microglia density from the same animal. There was a strong negative correlation between total microglia density and dying RGC density for either depletion strategy (Figure 4K). Overall, these findings suggest that CD11c-GFP+ microglia clear dying neurons, but are not selectively responsible or specialized to phagocytose apoptotic cells.

**Figure 4.**
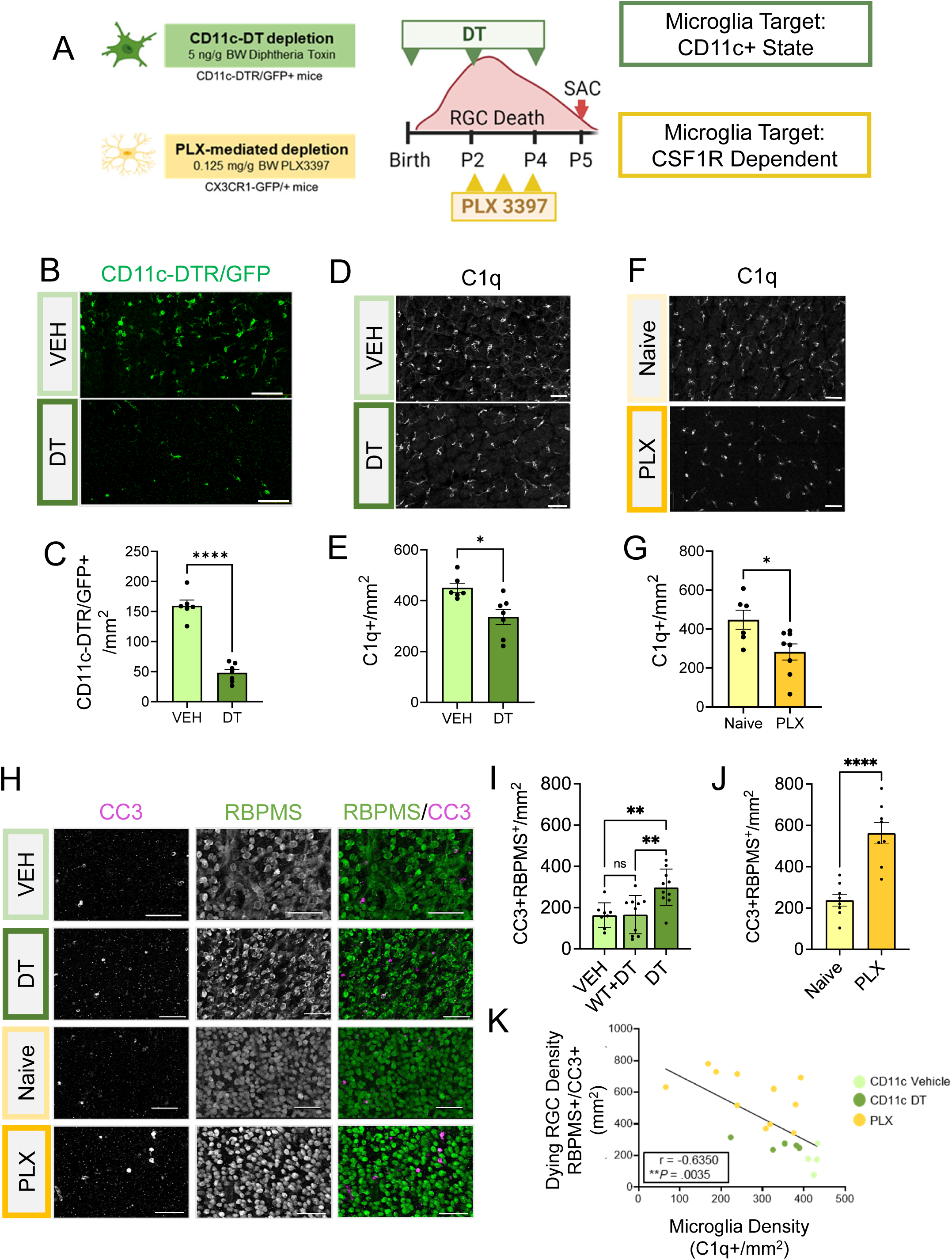
CD11c+ microglia are not solely responsible for phagocytic clearance of apoptotic neurons. (A) Depiction of two depletion strategies, Diphtheria toxin (DT) targeted ablation of CD11c-DTR/GFP+ microglia (Top) and PLX3397 (PLX)-mediated depletion to target more homeostatic microglia (Bottom). (B) Confocal images of CD11c-DTR/GFP+ microglia (green) in retinal flat mounts at P5 of Vehicle and DT-treated CD11c-DTR/GFP mice. Scale bar, 50μm. (C) Density of CD11c-DTR/GFP+ microglia in vehicle and DT-treated CD11c-DTR/GFP (n=6, n=7 animals) ± SEM. Unpaired t test \*\*\*\**P* <.0001. (D) Confocal images of C1q+ microglia (white) in immunostained retinal flat mounts at P5 of vehicle and DT-treated CD11c-DTR/GFP mice. Scale bar, 50μm. (E) Densities of C1q+ cells in retinas of vehicle and DT-treated CD11c-DTR/GFP mice (n=6, n=7 animals) ± SEM. Mann Whitney test *\*P =*.0140. (F) Max projected confocal images of C1q+ microglia in retinal flat mounts at P5 of vehicle and PLX-treated Cx3CR1/GFP+ mice. C1q (white). Scale bar, 50μm. (G) Density of C1q+ cells in retinas of naïve and PLX-treated CX3CR1-GFP (n=6, n=8) ± SEM. Unpaired t test *\*P =*.0239. (H) Max projected confocal images of apoptotic RGCs in retinal flat mounts at P5 of all conditions/genotypes. CC3 (magenta); RBPMS (green). Scale bar, 50μm. (I) Density of CC3+RBPMS+ cells in retinas from vehicle and DT-treated CD11c-DTR/GFP mice and DT-treated wildtype mice (WT) (n=8, n=10, n=10 animals respectively) ± SEM. Ordinary one-way ANOVA *\*\*P =*.0018 and Tukey’s multiple comparisons test: Veh vs DT \*\**P* =.006, WT+DT vs DT *\*\*P =*.0043, *NS*, not significant. (J) Density of CC3+RBPMS+ cells in naïve and PLX-treated CX3CR1-GFP/+ (n=8, n=8 animals) ± SEM. Unpaired t test \*\*\*\**P* <.0001 (K) Scatter plot of dying RGC density (CC3+RBPMS+/mm^2^) compared to microglial density (C1q+/mm^2^) of same animal (n=4 CD11c + vehicle, n=5 CD11c + DT, n=10 PLX animals). Pearson r= -0.6350 \*\**P* =.0035.

## Discussion

Single cell sequencing has revealed that microglia exist in diverse transcriptional states depending on local environmental cues. Less is known, however, regarding how dynamic and reversible these states are, the cues that produce specific states, and their relationship to cellular function. Here we address this by examining the emergence and regulation of CD11c+ microglia in developing retina. We first establish the recognition of apoptotic neurons as an important cue. CD11c+ microglia are linked to developmental waves of neuronal apoptosis, are more phagocytic and interact with apoptotic neurons, and expression of CD11c is modulated by signaling through the TAM receptor Axl, which can recognize phosphatidylserine exposure by apoptotic neurons (32). Second, using lineage tracing we provide evidence that microglia transition out of the CD11c+ state after the periods of neuronal apoptosis end, demonstrating that this state is reversible. Finally, we used selective ablation strategies to test whether CD11c+ microglia play a specialized role in apoptotic cell clearance and found that they are not selectively responsible, and that all microglia can participate in the elimination of apoptotic neurons. Altogether, this supports the notion that CD11c microglia in the retina arise from exposure to dying cells.

CD11c+ microglia have been reported in association with Aβ plaques, in developing, aged, and diseased white matter tracts of the brain, as well as the developing retina (8, 33). They have been implicated in development and repair of myelinated tracts (8-10), but their abundance in the developing retina, which lacks myelination, raises broader questions, including the environmental and molecular context for their emergence during development. In adult retina, CD11c+ microglia density is low, but following optic nerve crush, CD11c+ myeloid cells increase in number and play a prominent role in the clearance of retinal ganglion cell and axonal debris (34), with evidence that some CD11c+ cells are recruited into the retina from the myelinated optic nerve (35). The transcriptional profile of CD11c+ microglia is overlapping with the transcriptome signatures described for proliferative associate microglia (PAM) and axon tract associated microglia (ATM) in developing brain, and related to gene signatures associated with aging and neurodegenerative disease (8, 11, 14, 36, 37), raising the possibility of related cues driving these microglial states.

### CD11c+ microglia are driven by apoptosis

We found previously ∼50-60% of microglia within the developing postnatal retina express CD11c, and that this proportion was significantly reduced in Bax KO retina (6), which lack neuronal apoptosis (38). Here, we confirm that CD11c-GFP+ microglia density peaks postnatally in the retina and tracks with developmental apoptosis. We found that the majority of CD11c-GFP+ microglia interact with apoptotic cell bodies and have higher lysosomal content than their non-CD11c counterparts, consistent with enhanced phagocytosis. CD11c+ microglia have been linked to phagocytosis in multiple contexts. Within the developing brain, loss of “don’t-eat-me” signatures increases the number of CD11c+ microglia (39). While environmental cues driving this population in brain are unknown, consistent with our data, CD11c+ microglia in white matter tracts of the developing brain have been shown to engulf apoptotic oligodendrocytes and their precursors (4, 15). These data suggest microglial interactions with apoptotic cells may promote acquisition of the CD11c+ state. Consistent with this, our lineage analysis experiments found that CD11c-active microglia were more ameboid, had more phagocytic cups, and concentrated to layers undergoing developmental apoptosis whereas CD11c-lineage microglia had a branched morphology, less phagocytic cups, and distributed normally across all retinal layers. Despite this, we found that CD11c-GFP+ microglia were not any more efficient at the clearance of apoptotic neurons than their CD11c-negative counterparts. We found that the clearance of apoptotic neurons was directly proportional to the total number of microglia available, suggesting phagocytic clearance is not augmented in CD11c+ microglia.

### CD11c acquisition is regulated by Axl signaling

Our data and others show that the recognition of eat-me signatures is important for the acquisition of CD11c+ signature (5, 6, 9, 14, 29, 40). Conversely, “don’t eat-me” signature, Sirpα and CD47, were shown to constrain CD11c-expression through repression of eat-me signatures (39, 41). We found that early postnatal retina in Axl KO have a significant reduction in CD11c+ microglia, matching levels of CD11c+ microglia in Bax KO (6). We also found that microglial expression of Spp1 was reduced by Axl KO, consistent with other studies showing Spp1 expression by CD11c+ microglia (5, 14). In fact, CD11c+ microglia were reported to be the sole producer of Spp1 in the brain (14), and axon tract associated/CD11c+ microglia were shown to co-express Spp1 and were found to maintain structural integrity at cortical boundaries during fetal development (42). Taken together, our data implicate Axl in regulating the CD11c+ microglia state.

A previous study also implicated TAM receptors in the regulation of CD11c, with blockage of apoptotic neuron uptake through the inhibition of pan-Tam receptors and αVβ3 resulting in a marked reduction of CD11c-expression by microglia (43). However, we have previously shown that Axl is not required for the uptake and clearance of apoptotic neurons (5), suggesting that other Axl-mediated mechanisms may be involved. Interestingly, we also observed a marked increase in the proportion of microglia expression CD11c in CR3 KOs, suggesting a potential compensation by CD11c (CR4) for the loss of complement recognition by CD11b (CR3). CR3 is broadly expressed by microglia and is involved in the recognition of stressed or dying neurons in a variety of contexts, including the elimination of stressed RGCs embryonically and dying RGCs postnatally (5, 44). The observed increase in CD11c-expression with loss of CR3 expression may be attributed to compensatory upregulation, as both CR3 and CR4 are capable of recognizing deposition of iC3b, albeit CR4 is less readily positioned to bind iC3b under normal conditions (26, 27). Altogether, CD11c expression appears directly tied to the balance of eat-me cues versus don’t eat me cues, with a reduction of eat-me signaling through Axl translating into a reduction of CD11c+ microglia, and conversely, a reduction of don’t eat-me signals (like Sirpα and CD47) translating into an increase in CD11c+ microglia (39, 41).

### CD11c+ microglia are a transient and dynamically regulated state

With the proportion of CD11c + microglia in the retina declining as development progresses into adulthood, we asked whether CD11c+ microglia reintegrated back into the “homeostatic” pool of microglia. In brain, CD11c+ microglia are most abundant in cortical white matter tracts between postnatal days 3 through 8, with a sharp decline into adulthood (9, 10). Additionally, *in vitro* experiments show Cd11c-negative microglia, when cultured with apoptotic neurons, acquire CD11c-expression (14). These findings and fate-mapping experiments suggest CD11c+ microglia have the same origin as other brain microglia and do not arise from peripheral immune populations (10, 39). Here, we found that the proportion of CD11c-GFP+ microglia density dramatically shifts over time. Using a genetic lineage-tracing strategy, we find evidence that CD11c+ microglia reintegrate into the homeostatic pool of microglia within the retina, since from P5 to P30 the loss of CD11c-active microglia was proportional to the increase in CD11c-lineage microglia while the CD11c-negative microglia remain stable. To date, there have been conflicting data on whether CD11c+ microglia represent a stable subset that emerges in development and is maintained through adulthood (8, 9, 11), or a state achieved through recognition of molecular cue(s) (5, 6, 14). Altogether, our data suggest CD11c expression is transiently expressed in response to exposure to apoptotic neurons, and microglia shift back into a CD11c-state rather than being maintained as a stable subset.

## Materials

### Animal husbandry and procedures

Animals were treated within the University of Utah Institutional Animal Care and Use Committee (IACUC) guidelines and approval. Mice were housed in an AAALAC accredited animal facility with 12h light/12h dark cycles and ad libitum access to food and water. Both sexes were used for all experiments. PLX3397 (AdooQ BioScience,A15520) was dissolved in corn oil and 10% DMSO and administered to postnatal pups by intraperitoneal injection on P2-P4 at 0.125 mg/g body weight. Diphtheria Toxin (List Labs, #150) was dissolved in PBS and administered to postnatal pups by intraperitoneal injection at P0, P2, and P4 at 5ng/g of body weight.

### Mouse strains

The B6.129P2 (Cg)-*Cx3cr1^tm1Litt^*/J mice (22) were a gift from Richard Lang with permission from Steffen Jung. B6.FVB-*1700016L21Rik^Tg(Itgax-DTR/EGFP)57Lan^*/J (JAX 004509), C57BL/6J-Tg(Itgax-cre,-EGFP)4097Ach/J (JAX 007567), B6.Cg-*Gt(Rosa)26Sor^tm14(CAG-tdTomato)Hze^*/J (JAX 007914), and B6.129S4-Itgam^tm1Myd^/J (JAX 003991) were purchased from Jackson laboratories. The B6.129-Mertk^tm1Grl^/J and B6.129-Axl^tm1Grl^/J strains (25) were a gift from Dr. Greg Lemke.

### Tissue processing

Mice were decapitated following isoflurane asphyxiation, and whole heads were fixed in 4% PFA for 45 minutes to an hour. Heads were washed 3 times for 15 minutes in PBS and underwent 12–16-hour consecutive treatments with 15% and 30% sucrose in PBS at 4°C. Heads were then embedded in OCT compound (Tissue-Tek), stored at 80°C, and sectioned at 16μm or 30μm thickness. For retinal flat mounts, eyes were removed from the head and retinas were carefully dissected from the rest of the eye in ice cold 0.1M PBS, then washed in PBS for 10mins, and fixed in 4% PFA for 40mins at room temperature. For flow cytometry/FACS experiments, retinas were dissected RNAse-free conditions.

### Immunohistochemistry

Frozen sections were placed in ice-cold PBS for 10min, post-fixed for 10min in 4% PFA, washed in ice-cold PBS 3 times for 5min each, and then blocked for 1 hr at room temperature (0.2% triton-X, 10% BSA, 10% normal donkey serum in 0.01M PBS). Sections were incubated in primary antibody overnight at 4°C (0.2% triton-X, 5% BSA in 0.01M PBS). The following day, sections were washed 3 times with PBS for 5min each and incubated in secondary antibodies (5% BSA in 0.01M PBS) with Hoechst 33342 (1:5000, Thermo Fischer Scientific H3570) for 2 hrs at room temperature, washed, and mounted with Fluoroshield mounting medium with DAPI (Millipore Sigma, F6057). Flat mount immunostaining followed the same protocol except for primary antibody incubation, with flat mounts incubated in primary antibodies at 4°C for 2 days. Antibodies used include Goat polyclonal anti-GFP (1:2000, Abcam ab5450, RRID: AB_304897), Rabbit monoclonal anti-C1q (1:1500, Abcam an182451, RRID: AB_2732849), Rabbit polyclonal anti-IBA1 (1:1000, Wako 019-19741, RRID: AB_839504), Rabbit monoclonal anti-CC3 (1:500, BD Biosciences 559565, RRID: AB_397274), Guinea pig polyclonal anti-RBPMS (1:750, Millipore Sigma ABN1376, RRID: AB_2687403), Rat monoclonal anti-CD68 (1:250, Bio-Rad MCA1957, RRID: AB_322219), and Goat polyclonal anti-Osteopontin (1:100, Thermo Fischer PA5-34579, RRID: AB_2551931). Secondary antibodies were produced in donkey against goat, mouse, rabbit, guinea pig, and rat IgG, and conjugated to AlexaFluor -488, -555, or -647 (Thermo Fisher Scientific).

### Confocal microscopy

Confocal images were acquired on an inverted Nikon A1R Confocal Microscope, with images acquired at 20X objective and 3X digital zoom and 10% overlap multi-point stitching. Z plane stacks were at 0.8µm steps, with ∼13µm thickness to capture the NFL/GCL at 0.2mm pixel resolution. Flat mount retina images represent maximal-intensity projections of inner retina, primarily targeting GCL (NFL/GCL) from the central retina to the periphery, and acquisition settings were consistent across ages and genotypes.

### Fluorescence-activated cell sorting (FACS) and flow cytometry

2 retinas from an individual animal were pooled for each sample for Flow cytometry and FACS. Freshly dissected pure retinas were dissociated in PBS, 50 mM HEPES, 0.05 mg/ml DNase I (Sigma D4513), 0.025 mg/ml Liberase (Sigma 5401119001) for 35 min with intermediate trituration. Cells were passed through a 70 µm nylon cell strainer, washed with staining buffer (1X PBS, 2% BSA, 0.1% sodium azide, 0.05% EDTA), and red blood cells were lysed (eBioscience 00-4333-57). Cell counts were determined using a cell counter (Invitrogen Countess) and Fc block (BD Biosciences 553142) was added at 2 mL per 10^6^ cells. Antibodies used: BV421 Rat monoclonal anti-CD45 (BD Bioscience 563890, RRID: AB_2651151), AF 700 Rat monoclonal anti-CD11b (BD Bioscience 557960, RRID: AB_396960), PE-Cy7 Rat monoclonal anti-Ly6C (BD Bioscience 560593, RRID: AB_1727557), APC Hamster monoclonal anti-CD11c (BD Bioscience Pharmingen 561119, RRID: AB_10562405), and APC Rat monoclonal CD206 (Thermo Fisher Scientific 141708, RRID: AB_10896057) were applied for 30 min on ice. Cells were washed, pelleted, and resuspended in 500 µL staining buffer. FACS was performed using a BD FACS Aria cell sorter at the University of Utah Flow Cytometry Core. Forward and side scatter were used to eliminate debris and both the width and area of the forward and side scatter was used to discriminate singlets. ∼1 million singlet events were recorded for analysis, and gates for flow analysis were determined using FMO control. Analysis was conducted using FlowJo software (Flowjo, LLC, Ashland, Oregon).

### CD11c-lineage bulk-seq

Bulk RNA-sequencing was performed on FACs sorted cells from CD11c-Cre-gfp;Rosa-TDTom animals at P5. One sample each of Cd11c lineage cells (GFP-TdTom+, 5441 cells) and Cd11c active cells (GFP+TdTom+, 4041 cells) were sorted from 8 retinas (4 female and 4 male). Total RNA was prepared with the RNeasy Plus Kit (Qiagen 74034) and RNA Clean and Concentrator kit (Zymo R1013) and hybridized with NEBNext rRNA Depletion Kit v2 (NEB E7400) to substantially diminish rRNA from the samples. Stranded RNA sequencing libraries were prepared as described using the NEBNext Ultra II Directional RNA Library Prep Kit for Illumina (NEB E7760L). Purified libraries were qualified on an Agilent Technologies 4150 TapeStation using a D1000 ScreenTape assay (5067-5582 and 5067-5583). The molarity of adapter-modified molecules was defined by quantitative PCR using the Kapa Biosystems Kapa Library Quant Kit (KK4824). Individual libraries were normalized to 5 nM in preparation for Illumina sequence analysis. Libraries were sequenced on an Illumina NovaSeq 6000 S4 with reagent kit v1.5 to produce 150x150 bp paired-end sequences to a depth of greater than 50 million paired reads. Illumina adapters were trimmed from reads with cutadapt v3.5, trimming after 6 matching bases and discarding trimmed reads shorter than 20 bases. Trimmed reads were aligned with STAR v2.7.9a in two pass mode to indexed genome GRCm39, created with splice junctions from Ensembl release 104 and a maximum overhang of 124 bases, to generate a BAM alignment file sorted by coordinates. Uniquely aligned fragment counts were assigned to Ensembl 104 transcripts from the alignment files with featureCounts v1.6.3.edgeR v4.0.16 was used to calculate log fold changes and log CPM from read counts with the exactTest function. Raw and processed data files are available at the Gene Expression Omnibnus under GEO accession number GSE269549.;

### Image analysis

All counts were blinded and manually performed using Nikon Elements software (Melville, NY). For double positive CC3+RBPMS+ or single RBPMS+ counts of retinal flat mounts, two ROIs of roughly 0.4 mm^2^ of central and periphery of dorsal retina were analyzed. Images were maximal-intensity projected and roughly 13µm thick, spanning the NFL to the GCL. For quantification of CD11c-GTFP+ microglia in retinal flat mounts, the entire retina, ranging from 1.5 to 8 mm^2^, was analyzed across all retinal depths, ranging from 25 to 55µm. Microglial proportion was determined as CD11c-GFP+/total microglia (C1q+). CD11c-GFP+ localization analysis was performed on entire cross sections of 3 retinas. For 3D reconstructions of microglia-apoptotic body interactions, IMARIS software (Bit-plane) was used. In brief, high resolution confocal images (20X objective, 8X digital zoom) through Z (0.16µm steps) were uploaded into IMARIS to create 3D renderings. For CD68 and C1q area analysis, P3 retinal flat mount imaging and analysis spanned ∼13μm spanning the NFL to GCL only for the dorsal quadrant of the right eye of each animal analyzed. The lysosomal content (CD68 area) was analyzed against total microglia area (union of C1q and CD68 channels), with quantification of CD68/C1q+CD68 in CD11c-GFP+ versus CD11c-GFP-microglia. For phagocytic cup analysis, a “phagocytic cup” was determined based upon a cup-shaped invagination of the plasma membrane formed around cellular debris or dying/dead cells located at the tip of microglial processes. Microglia phagocytic cups were counted in P5 retinal flat mounts stained with CD68 in CD11c-Cre^GFP/+^; Rosa^tdTomato/+^, focusing on CD11c-active (GFP+tdTomato+) and CD11c-lineage (GFP-tdTomato+) microglia.

### Statistical methods

All image and flow data were analyzed using Prism 10 software (GraphPad, La Jolla, CA). We first tested for normality using four tests: Anderson-Darling, D’Agostino & Pearson, Shapiro-Wilk, and Kolmogorov-Smirnov test. Failure of any one test resulted in non-parametric tests, Mann-Whitney for pair-wise comparisons and Kruskal-Wallis for comparisons of more than three samples. We tested for heteroscedasticity using an F-test for pairwise comparisons and Brown-Forsythe test for comparisons of three or more. If not significantly different, we either ran a two-tailed T test, Ordinary one-way ANOVA with posthoc Tukey’s multiple comparison test, or Ordinary two-way ANOVA with Sidak’s multiple comparison test. If standard deviations were significantly different between groups, we ran a Welch’s ANOVA with posthoc Dunnett’s T3 multiple comparison’s test. For all data that is presented as the mean, error bars indicate the standard error of the mean, SEM. We used a 95% confidence interval and a *P* value of < .05 was set for rejecting the null hypothesis.

## Supporting information

**Supplementary Figure S1.**
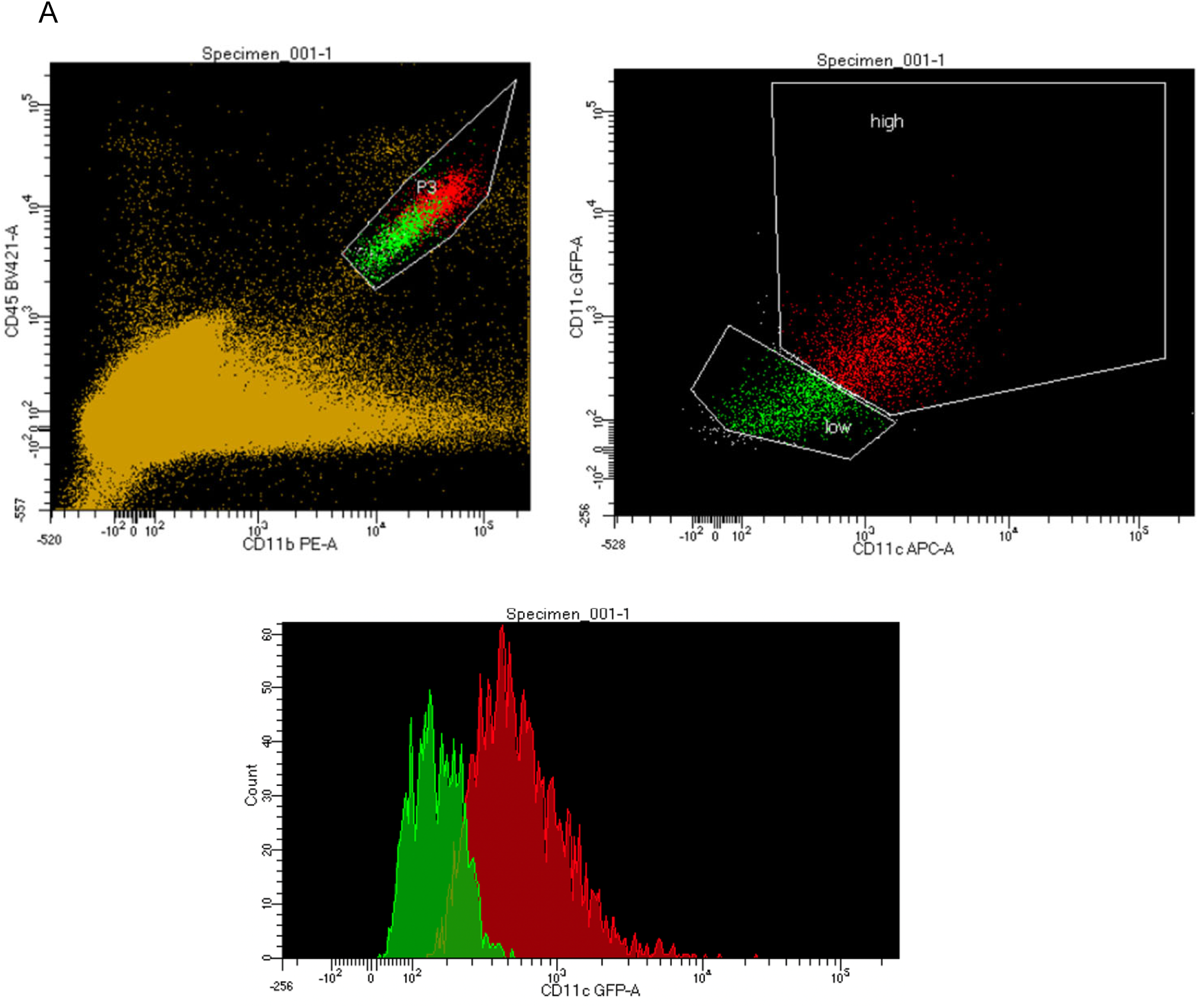
CD11c-GFP correlates with microglia surface expression of CD11c. Related to. **Figure 1**. (A) (left) Gating strategy for FACS of CD45+CD11b+ microglia and (right) CD11c-GFP expression with CD11c-antibody (APC). (bottom) Histogram of CD11c+ and CD11c-populations along CD11c-GFP+ signal. Left and bottom back colored based on gating on the right.

**Supplementary Figure S2.**
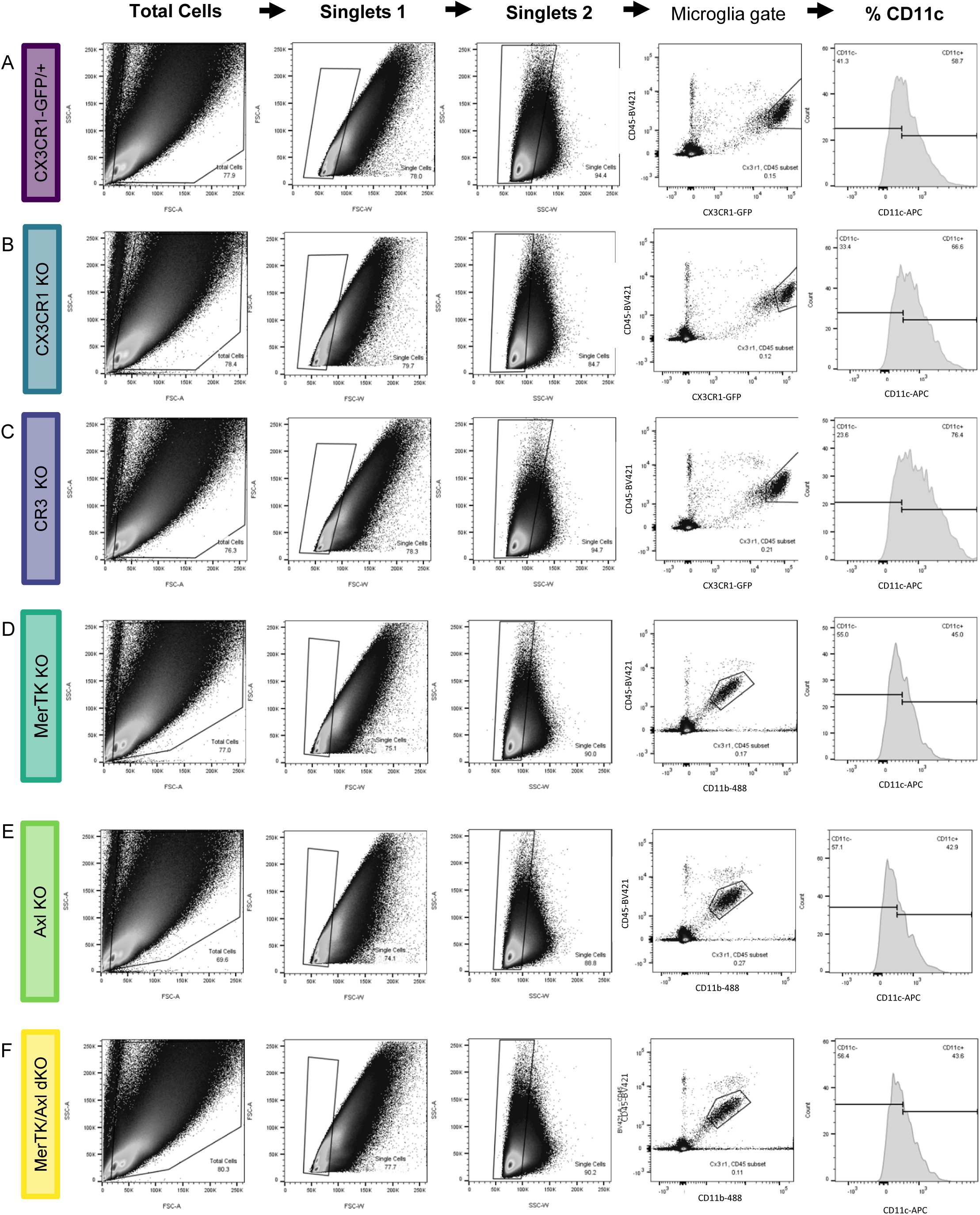
Flow cytometry analysis of CD11c^Hi^ microglia with loss of candidate receptors. Related to. **Figure 3**. (A – F) Representative flow plots illustrating the proportion of CD45+GFP+ or CD45+CX3CR1-GFP+ microglia that are CD11c^Hi^. ≥ 2 litters collected for each genotype, with (A) CX3CXR1-GFP/+ n=10 (B) CX3CR1 KO n=6, (C) CR3 KO n=6, (D) MerTK KO n=9, (E) Axl KO n=8, and (F) MerTK/Axl dKO n=6 animals.

**Supplementary Figure S3.**
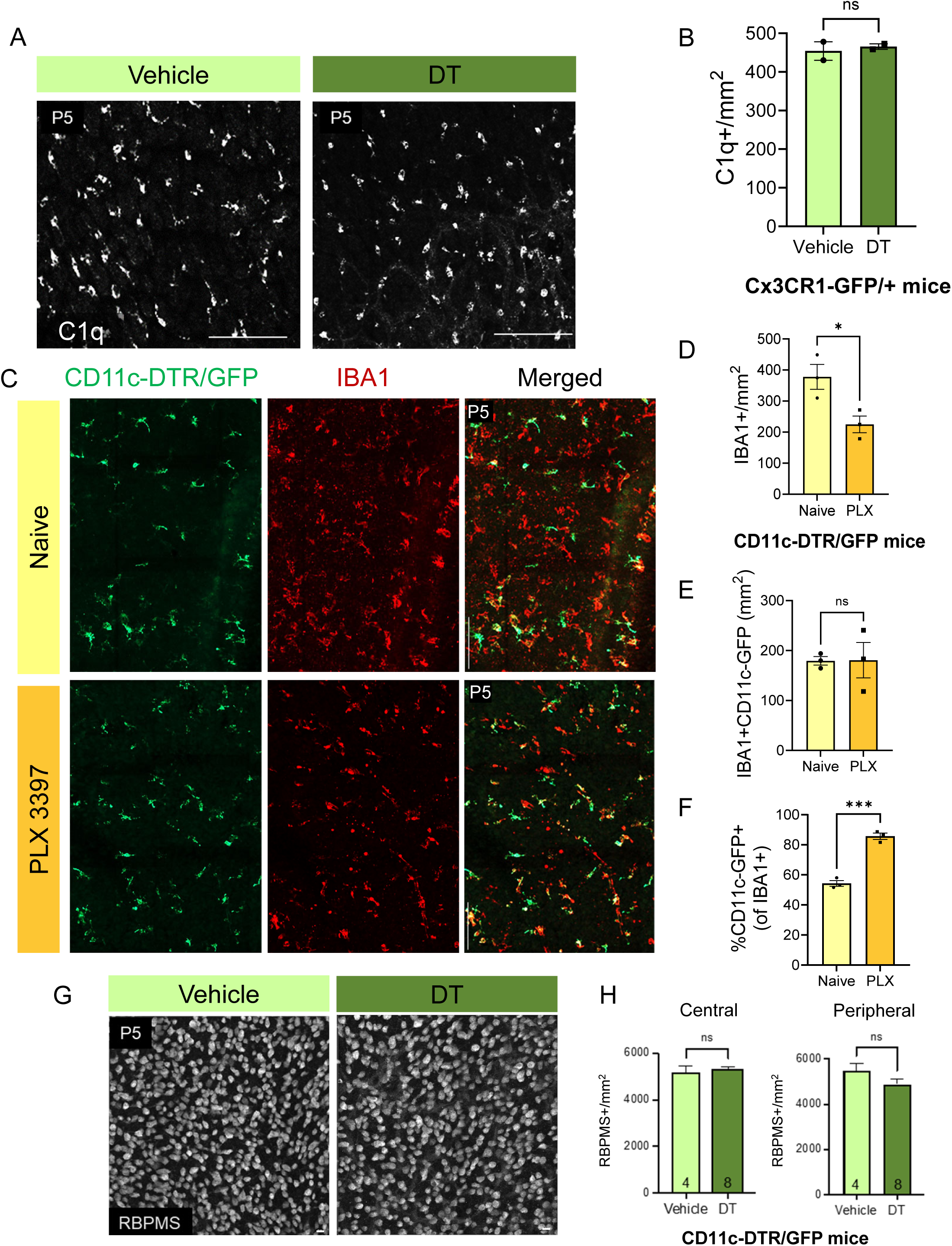
Diphtheria toxin administration does not affect microglia without primate DTR expression, PLX administration enriches for CD11c-GFP+ microglia. Related to Figure 4. (A) Confocal images of microglia (C1q+) in retinal flat mounts at P5 of Vehicle and DT-treated CX3CR1-GFP/+ mice. C1q+ (white). Scale bar, 50μm. (B) Density of C1q+ microglia in vehicle and DT-treated CX3CR1-GFP/+ mice at P5 (n=2 animals) ± SEM. Unpaired t test *NS,* not significant. (C) Confocal retinal flat mount images from naïve and PLX treated CD11c-DTR/GFP+ mice at P5. CD11c-DTR/GFP microglia (GFP+IBA1+) and total microglia (IBA1+, red) Scale bar, 50μm. (D – F) Densities of naïve and PLX treated CD11c-DTR/GFP+ mice, with (D) total microglia density (IBA1+; n=3, n=3 animals), (E) CD11c-DTR/GFP+ microglia density (IBA1+CD11c-DTR/GFP+; n=3, n=3 animals), and (F) proportion of microglia (IBA1+) expressing CD11c-DTR/GFP (n=3, n=3 animals) ± SEM. Unpaired t test \*\*\**P* = .0004, \**P =* .0340, *NS*, not significant. (G) Confocal images of RGCs (RBPMS+) in retinal flat mounts at P5 of vehicle and DT treated CD11c-DTRG/GFP+. RBPMS (white). Scale bar, 50μm. (H) Densities of vehicle and DT-treated CD11c-DTR/GFP RGCs in (Left) central region and (Right) peripheral region of dorsal leaf (vehicle n=4, DT n=8 animals) ± SEM. Unpaired t test *NS,* not significant, Scale bars, 50μm.

## Acknowledgments

Cartoons were created with BioRender. We thank the University of Utah Office of Comparative Medicine for animal husbandry. Research reported in the publication utilized the University of Utah Flow Cytometry Core at Huntsman Cancer Institute at the University of Utah, which is supported by the Office of the Director of the NIH and NCI. Research reported in this publication utilized the High-Throughput Genomics and Bioinformatic Analysis Shared Resource at the Huntsman Cancer Institute at the University of Utah. The support and resources from the Center for High Performance Computing at the University of Utah are gratefully acknowledged. This content is solely the responsibility of the authors and does not necessarily represent the official views of the NIH.

## Author contributions

### Conceptualization

Nathaniel Ghena, Sarah R Anderson, Monica L Vetter

### Formal analysis

Nathaniel Ghena, Sarah R Anderson, Jacqueline M Roberts, Emmalyn Irvin, Joon Schwakopf, Alejandra Bosco

### Funding acquisition

Nathaniel Ghena, Monica L Vetter

### Investigation

Nathaniel Ghena, Sarah R Anderson, Jacqueline M Roberts, Monica L Vetter

### Methodology

Nathaniel Ghena, Sarah R Anderson, Jacqueline M Roberts, Emmalyn Irvin, Joon Schwakopf, Alejandra Bosco, Monica L Vetter

### Resources

Nathaniel Ghena, Sarah R Anderson, Joon Schwakopf, Emmalyn Irvin, Monica L Vetter

### Visualization

Nathaniel Ghena, Sarah R Anderson, Jacqueline M Roberts

### Writing – original draft

Nathaniel Ghena, Sarah R Anderson, Monica L Vetter

### Writing – review & editing

Nathaniel Ghena, Sarah R Anderson, Jacqueline M Roberts, Emmalyn Irvin, Joon Schwakopf, Alejandra Bosco, Monica L Vetter

## References

1. Paolicelli RC, Sierra A, Stevens B, Tremblay ME, Aguzzi A, Ajami B, et al. Microglia states and nomenclature: A field at its crossroads. Neuron. 2022;110(21):3458–83. 10.1016/j.neuron.2022.10.020

2. Prinz M, Masuda T, Wheeler MA, Quintana FJ. Microglia and Central Nervous System-Associated Macrophages-From Origin to Disease Modulation. Annu Rev Immunol. 2021;39:251–77. 10.1146/annurev-immunol-093019-110159

3. Hammond TR, Dufort C, Dissing-Olesen L, Giera S, Young A, Wysoker A, et al. Single-Cell RNA Sequencing of Microglia throughout the Mouse Lifespan and in the Injured Brain Reveals Complex Cell-State Changes. Immunity. 2019;50(1):253–71 e6. 10.1016/j.immuni.2018.11.004

4. Li Q, Cheng Z, Zhou L, Darmanis S, Neff NF, Okamoto J, et al. Developmental Heterogeneity of Microglia and Brain Myeloid Cells Revealed by Deep Single-Cell RNA Sequencing. Neuron. 2019;101(2):207–23 e10. 10.1016/j.neuron.2018.12.006

5. Anderson SR, Roberts JM, Ghena N, Irvin EA, Schwakopf J, Cooperstein IB, et al. Neuronal apoptosis drives remodeling states of microglia and shifts in survival pathway dependence. Elife. 2022;11 10.7554/eLife.76564

6. Anderson SR, Roberts JM, Zhang J, Steele MR, Romero CO, Bosco A, et al. Developmental Apoptosis Promotes a Disease-Related Gene Signature and Independence from CSF1R Signaling in Retinal Microglia. Cell Rep. 2019;27(7):2002–13 e5. 10.1016/j.celrep.2019.04.062

7. Pitts KM, Margeta MA. Myeloid masquerade: Microglial transcriptional signatures in retinal development and disease. Front Cell Neurosci. 2023;17:1106547. 10.3389/fncel.2023.1106547

8. Benmamar-Badel A, Owens T, Wlodarczyk A. Protective Microglial Subset in Development, Aging, and Disease: Lessons From Transcriptomic Studies. Front Immunol. 2020;11:430. 10.3389/fimmu.2020.00430

9. Wlodarczyk A, Holtman IR, Krueger M, Yogev N, Bruttger J, Khorooshi R, et al. A novel microglial subset plays a key role in myelinogenesis in developing brain. EMBO J. 2017;36(22):3292–308. 10.15252/embj.201696056

10. Hagemeyer N, Hanft KM, Akriditou MA, Unger N, Park ES, Stanley ER, et al. Microglia contribute to normal myelinogenesis and to oligodendrocyte progenitor maintenance during adulthood. Acta Neuropathol. 2017;134(3):441–58. 10.1007/s00401-017-1747-1

11. Nomaki K, Fujikawa R, Masuda T, Tsuda M. Spatiotemporal dynamics of the CD11c(+) microglial population in the mouse brain and spinal cord from developmental to adult stages. Mol Brain. 2024;17(1):24. 10.1186/s13041-024-01098-2

12. Kamphuis W, Kooijman L, Schetters S, Orre M, Hol EM. Transcriptional profiling of CD11c-positive microglia accumulating around amyloid plaques in a mouse model for Alzheimer’s disease. Biochim Biophys Acta. 2016;1862(10):1847–60. 10.1016/j.bbadis.2016.07.007

13. Keren-Shaul H, Spinrad A, Weiner A, Matcovitch-Natan O, Dvir-Szternfeld R, Ulland TK, et al. A Unique Microglia Type Associated with Restricting Development of Alzheimer’s Disease. Cell. 2017;169(7):1276–90 e17. 10.1016/j.cell.2017.05.018

14. Shen X, Qiu Y, Wight AE, Kim HJ, Cantor H. Definition of a mouse microglial subset that regulates neuronal development and proinflammatory responses in the brain. Proc Natl Acad Sci U S A 2022;119(8) 10.1073/pnas.2116241119

14. Nemes-Baran AD, White DR, DeSilva TM. Fractalkine-Dependent Microglial Pruning of Viable Oligodendrocyte Progenitor Cells Regulates Myelination. Cell Rep. 2020;32(7):108047. 10.1016/j.celrep.2020.108047

15. Jung S, Unutmaz D, Wong P, Sano G-I, De los Santos K, Sparwasser T, et al. In Vivo Depletion of CD11c+ CD8+ Dendritic Cells Abrogates Priming of T Cells by Exogenous Cell-Associated Antigens. Immunity. 2002;17(2):211–20.

16. Young RW. Cell Death During Differentiation of the Retina in the Mouse. Journal of Comparative Neurology. 1984;229:362–73.

17. Santos AM, Calvente R, Tassi M, Carrasco MC, Martin-Oliva D, Marin-Teva JL, et al. Embryonic and postnatal development of microglial cells in the mouse retina. J Comp Neurol. 2008;506(2):224–39. 10.1002/cne.21538

18. Stranges PB, Watson J, Cooper CJ, Choisy-Rossi CM, Stonebraker AC, Beighton RA, et al. Elimination of antigen-presenting cells and autoreactive T cells by Fas contributes to prevention of autoimmunity. Immunity. 2007;26(5):629–41. 10.1016/j.immuni.2007.03.016

19. Madisen L, Zwingman TA, Sunkin SM, Oh SW, Zariwala HA, Gu H, et al. A robust and high-throughput Cre reporting and characterization system for the whole mouse brain. Nat Neurosci. 2010;13(1):133–40. 10.1038/nn.2467

20. Krasemann S, Madore C, Cialic R, Baufeld C, Calcagno N, El Fatimy R, et al. The TREM2-APOE Pathway Drives the Transcriptional Phenotype of Dysfunctional Microglia in Neurodegenerative Diseases. Immunity. 2017;47(3):566–81 e9. 10.1016/j.immuni.2017.08.008

21. Jung S, Aliberti J, Graemmel P, Sunshine MJ, Kreutzberg GW, Sher A, et al. Analysis of fractalkine receptor CX(3)CR1 function by targeted deletion and green fluorescent protein reporter gene insertion. Mol Cell Biol. 2000;20(11):4106–1414. doi: 10.1128/MCB.20.11.4106-4114.2000

22. Coxon A, Rieu P, Barkalow FJ, Askari S, Sharpe AH, von Andrian UH, et al. A Novel Role for the beta-2 Integrin CD11b/CD18 in Neutrophil Apoptosis: A Homeostatic Mechanism in Inflammation. Immunity. 1996;5:653–66.

23. Fourgeaud L, Traves PG, Tufail Y, Leal-Bailey H, Lew ED, Burrola PG, et al. TAM receptors regulate multiple features of microglial physiology. Nature. 2016;532(7598):240–4. 10.1038/nature17630

24. Lu Q, Gore M, Zhang Q, Camenisch T, Boast S, Casagranda F, et al. Tyro-3 family receptors are essential regulators of mammalian spermatogenesis. Nature. 1999;398(6729):723-8. doi: 10.1038/19554

25. Sandor N, Lukacsi S, Ungai-Salanki R, Orgovan N, Szabo B, Horvath R, et al. CD11c/CD18 Dominates Adhesion of Human Monocytes, Macrophages and Dendritic Cells over CD11b/CD18. PLoS One. 2016;11(9):e0163120. 10.1371/journal.pone.0163120

26. Ross GD, Reed W, Dalzell JG, Becker SE, Hogg N. Macrophage cytoskeleton association with CR3 and CR4 regulates receptor mobility and phagocytosis of iC3b-opson ized erythrocytes. Journal of Leukocyte Biology. 1992;51:109–17.

27. Probst HC, Tschannen K, Odermatt B, Schwendener R, Zinkernagel RM, Van Den Broek M. Histological analysis of CD11c-DTR/GFP mice after in vivo depletion of dendritic cells. Clin Exp Immunol. 2005;141(3):398–404. 10.1111/j.1365-2249.2005.02868.x

29. A. Qiu Y, Shen X, Ravid O, Atrakchi D, Rand D, Wight AE, et al. Definition of the contribution of an Osteopontin-producing CD11c(+) microglial subset to Alzheimer’s disease. Proc Natl Acad Sci U S A 2023;120(6):e2218915120. 10.1073/pnas.2218915120

28. Pappenheimer AM, Harper AA, Moynihan M, Brockes JP. Diphtheria Toxin and Related Proteins: Effect of Route of Injection on Toxicity and the Determination of Cytotoxicity for Various Cultured Cells. The Journal of Infectious Disease. 1982;145:94–102.

29. Elmore MR, Najafi AR, Koike MA, Dagher NN, Spangenberg EE, Rice RA, et al. Colony-stimulating factor 1 receptor signaling is necessary for microglia viability, unmasking a microglia progenitor cell in the adult brain. Neuron. 2014;82(2):380–97. 10.1016/j.neuron.2014.02.040

30. Lemke G. Phosphatidylserine Is the Signal for TAM Receptors and Their Ligands. Trends Biochem Sci. 2017;42(9):738–48. 10.1016/j.tibs.2017.06.004

31. Anderson SR, Vetter ML. Developmental roles of microglia: A window into mechanisms of disease. Dev Dyn. 2019;248(1):98–117. 10.1002/dvdy.1

32. Lehmann U, Heuss ND, McPherson SW, Roehrich H, Gregerson DS. Dendritic cells are early responders to retinal injury. Neurobiol Dis. 2010;40(1):177–84. 10.1016/j.nbd.2010.05.022

33. Heuss ND, Pierson MJ, Roehrich H, McPherson SW, Gram AL, Li L, et al. Optic nerve as a source of activated retinal microglia post-injury. Acta Neuropathol Commun. 2018;6(1):66. 10.1186/s40478-018-0571-8

34. Holtman IR, Raj DD, Miller JA, Schaafsma W, Yin Z, Brouwer N, et al. Induction of a common microglia gene expression signature by aging and neurodegenerative conditions: a co-expression meta-analysis. Acta Neuropathol Commun. 2015;3:31. 10.1186/s40478-015-0203-5

35. Holtman IR, Skola D, Glass CK. Transcriptional control of microglia phenotypes in health and disease. J Clin Invest. 2017;127(9):3220–9. 10.1172/JCI90604

36. Pequignot MO, Provost AC, Salle S, Taupin P, Sainton KM, Marchant D, et al. Major role of BAX in apoptosis during retinal development and in establishment of a functional postnatal retina. Dev Dyn. 2003;228(2):231–8. 10.1002/dvdy.10376

37. Sato-Hashimoto M, Nozu T, Toriba R, Horikoshi A, Akaike M, Kawamoto K, et al. Microglial SIRPalpha regulates the emergence of CD11c(+) microglia and demyelination damage in white matter. Elife. 2019;8 10.7554/eLife.42025

38. Kohno K, Shiraska R, Yoshihara K, Mikuriya S, Tanaka K, Takanami K, et al. A spinal microglia population invovled in remitting and relapsing neuropathic pain. Science. 2022;376:86–90. DOI:10.1126/science.abf6805

41. A. Wu J, Wu H, An J, Ballantyne CM, Cyster JG. Critical role of integrin CD11c in splenic dendritic cell capture of missing-self CD47 cells to induce adaptive immunity. Proc Natl Acad Sci U S A 2018;115(26):6786–91. 10.1073/pnas.1805542115

39. Lawrence AR, Canzi A, Bridlance C, Olivie N, Lansonneur C, Catale C, et al. Microglia maintain structural integrity during fetal brain morphogenesis. Cell. 2024;187(4):962–80 e19. 10.1016/j.cell.2024.01.012

40. Zagorska A, Traves PG, Lew ED, Dransfield I, Lemke G. Diversification of TAM receptor tyrosine kinase function. Nat Immunol. 2014;15(10):920–8. 10.1038/ni.2986

41. Anderson SR, Zhang J, Steele MR, Romero CO, Kautzman AG, Schafer DP, et al. Complement Targets Newborn Retinal Ganglion Cells for Phagocytic Elimination by Microglia. J Neurosci. 2019;39(11):2025–40. 10.1523/JNEUROSCI.1854-18.2018

